# A zinc-finger fusion protein refines Gal4-defined neural circuits

**DOI:** 10.1101/228718

**Authors:** Shamprasad Varija Raghu, Farhan Mohammad, Chua Jia Yi, Claudia S. Barros, Joanne Lam, Mavis Loberas, Sadhna Sahani, Adam Claridge-Chang

## Abstract

The analysis of behavior requires that the underlying neuronal circuits are identified and genetically isolated. In several major model species—most notably *Drosophila*, neurogeneticists identify and isolate neural circuits with a binary heterologous expression-control system: Gal4–UASG. One limitation of Gal4–UASG is that expression patterns are often too broad to map circuits precisely. To help refine the range of Gal4 lines, we developed an intersectional genetic AND operator. Interoperable with Gal4, the new system’s key component is a fusion protein in which the DNA-binding domain of Gal4 has been replaced with a zinc finger domain with a different DNA-binding specificity. In combination with its cognate binding site (UASZ) the <u>z</u>inc-finger-replaced G<u>a</u>l4 (‘Zal1’) was functional as a standalone transcription factor. Zal1 transgenes also refined Gal4 expression ranges when combined with UASGZ, a hybrid upstream activation sequence. In this way, combining Gal4 and Zal1 drivers captured restricted cell sets compared with single drivers and improved genetic fidelity. This intersectional genetic AND operation presumably derives from the action of a heterodimeric transcription factor: Gal4-Zal1. Configurations of Zal1–UASZ and Zal1-Gal4-UASGZ are versatile tools for defining, refining, and manipulating targeted neural expression patterns with precision.

## Introduction

For the analysis of neural circuits and behavior, neuroscientists use transgenic techniques to isolate neuronal groups with precision. Neurogeneticists working with the vinegar fly *Drosophila melanogaster* have developed a sophisticated, versatile toolkit that includes a foundational transcriptional system for mapping and manipulating neural circuits: Gal4–UASG (Brand & Perrimon 1993). This system typically uses two fusion transgenes: endogenous fly enhancer sequences are placed upstream of the yeast transcription factor Gal4; effector transgenes are fused to Gal4’s upstream activation sequence (UASG). This arrangement places the effector under the *in trans* transcriptional control of the enhancer (Brand & Perrimon 1993). The Gal4–UASG method has been used for cell-specific genetic rescue, gene overexpression, reporter expression, RNA-interference screens, optogenetic physiology, and many other applications (Bellen et al. 2010; Mohammad et al. 2017). While this tool is vitally useful, one challenge to dissecting neuron–behavior relationships has been that Gal4-linked enhancers often capture more cells than are functionally relevant. To improve the precision of transgene expression, neural circuit analysis uses a variety of molecular strategies to produce AND and NOT genetic logic, producing expression refinements by intersection. Intersectional methods use either a repressor of Gal4, a targeted recombinase system, a leucine-zipped split-Gal4, or a combination. The native Gal4 repressor, Gal80, is used as a genetic NOT operator to exclude expression from a subset of cells captured by a driver (Suster et al. 2004). The flippase (Flp) recombinase specifically excises genomic sequences flanked by flippase recognition target (FRT) sites. In the Flp-out method, Flp is transiently expressed under the control of a heat shock promoter to both generate AND and NOT operations (Wong et al. 2002). Stochastic single-cell specificity can be achieved with the ‘mosaic analysis with repressible cell marker’ (MARCM) technique (Lee & Luo 1999). Flp-FRT is also used in the ‘Flippase-induced intersectional Gal80/Gal4 repression’ (FINGR) intersectional method (Bohm et al. 2010), wherein stable, elevated levels of Flp are expressed from an enhancer to add or remove Gal80 expression from a subset of Gal4 driver cells with some stochasticity (Sivanantharajah & Zhang 2015). The split-Gal4 method uses a bipartite Gal4 variant, in which a heterodimerization leucine zipper joins the DNA-binding and activation domains; it is active as a transcription factor when both components are expressed in the same cell, producing AND logic between the two half-drivers (Luan et al. 2006). A non-intersectional approach to improving cell set specificity uses driver lines constructed with small enhancer fragments instead of large upstream regions (Pfeiffer et al. 2008; Jenett et al. 2012; Kvon et al. 2014). Such genomic fragments contain fewer enhancer modules, so they tend to express in more restricted anatomical ranges: an estimated 4- to 10-fold greater specificity compared with enhancer traps (Pfeiffer et al. 2008).

In light of the extensive Gal4 resources currently available, we aimed to develop an tool that would refine existing Gal4 lines. The DNA-binding domain of Gal4 is a zinc finger that can be substituted with another domain, conferring novel DNA-binding affinity *in vitro* (Pomerantz et al. 1998). We implemented and tested a zinc finger variant of Gal4 that works both as a standalone binary transcription system and as a genetic AND operator in combination with existing Gal4 lines. Using several enhancer sequences associated with particular neurotransmitter systems, we demonstrated that the variant transcription factor—termed <u>Z</u>inc finger-replaced G<u>a</u>l4 (Zal1)—can drive expression from a corresponding upstream activating sequence, termed UASZ. When co-expressed in the same cells, Gal4 and Zal1 were active in the presence of a hybrid upstream activation sequence that contained asymmetric binding sites (UASGZ) for the Gal4-Zal1 heterodimer. This method allowed targeting of expression to neurons in which both transcription factor types are expressed. The Zal1-Gal4-UASGZ system will enable the refinement of existing Gal4 lines to isolate precise neuronal types.

## Results

### Ternary UAS expression system design

Gal4 binds to its cognate upstream activating DNA motif, referred to here as UASG (Figure 1A). Gal4 can be used to drive specific expression of a responder transgene (e.g. green fluorescent protein, GFP) in defined cell types such as specific *Drosophila* neurons (Figure 1B). Pomerantz and colleagues previously designed a transcription-factor fragment that fused the first two zinc fingers of mouse transcription activator EGR1 (previously referred to as ZIF268) with the linker and dimerization domains of Gal4 (Pomerantz et al. 1998). In an *in vitro* study, they showed that the resulting truncated fusion protein, <u>z</u>inc <u>f</u>inger <u>G</u>al4 <u>d</u>imerization <u>1</u> (ZFGD1), bound to DNA containing its corresponding UAS (here termed UASZ), a palindromic site with inverted EGR1 finger-binding sites (Figure 1C). Using the same fusion design as ZFGD1, we generated a gene encoding a full-length transcription factor, <u>z</u>inc finger-replaced G<u>a</u>l4 (Zal1), to be used *in vivo* to activate genes placed downstream of a UASZ tandem repeat (Figure 1D). Since heterodimeric ZFGD1 proteins assemble *in vitro* and specifically bind to hybrid UAS sites in DNA (Pomerantz et al. 1998), we anticipated that a full-length heterodimeric Gal4/Zal1 transcription factor would form *in vivo,* bind hybrid sites in the genome (Figure 1E), and activate a UASGZ-controlled responder in cells where Gal4 and Zal1 are co-expressed (Figure 1F).

**Figure 1.**
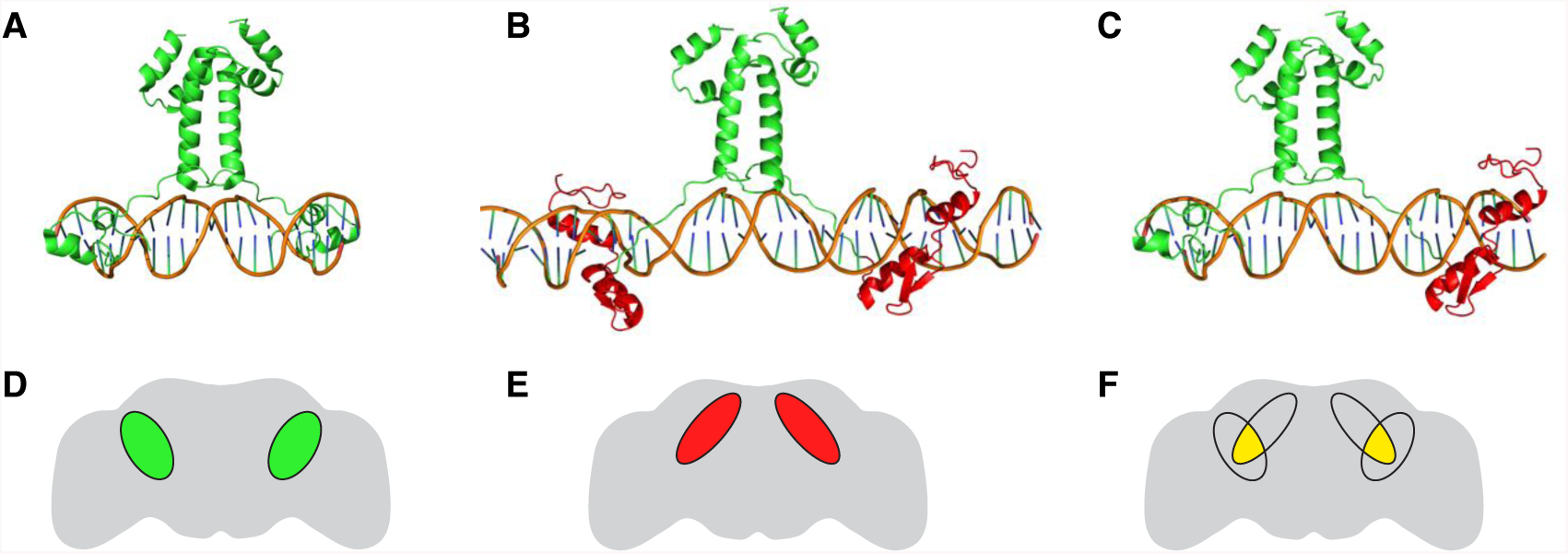
Structural models of Gal4 and Zal1; the experimental expression concept. **A.** Structural model of Gal4 protein domains in the native homodimeric configuration; two zinc fingers constitute the DNA-binding domain. Only the DNA-binding domain, linker and dimerization domain of Gal4 are shown. **B.** A hypothetical expression pattern for Gal4 homodimer driving expression from a UASG effector gene in the adult fly brain. **C.** A hypothetical structural model of Zal1 protein in which the zinc fingers of Gal4 are replaced with fingers 1 and 2 from the crystal structure of EGR1, shown in red. **D.** A hypothetical expression pattern for Zal1 homodimer driving expression from a UASZ effector gene. **E.** A model of the Gal4-Zal1 heterodimer. **F.** A hypothetical expression pattern produced by Gal4-Zal1 heterodimer in the presence of a UASGZ effector gene.

### *VGlut-Zal1* drives broad *UASZ-GFP* expression

Transgenic flies were prepared to carry *Zal1* fused to the *vesicular glutamate transporter (VGlut)* enhancer (Daniels et al. 2008). Progeny of *VGlut-Zal1* crossed with *UASZ-GFP* expressed GFP throughout the brain (Figure 2A–B). The *VGlut-Zal1* pattern differed from that of *VGlut-Gal4* (Figure 2C–D), possibly because *VGlut* enhancer fusions capture non-comprehensive cell sets (Daniels et al. 2008) and the expression variation that can arise from genomic insertion sites and vector design (Ni et al. 2008). This result demonstrates that Zal1 is functional in the *Drosophila* brain.

**Figure 2.**
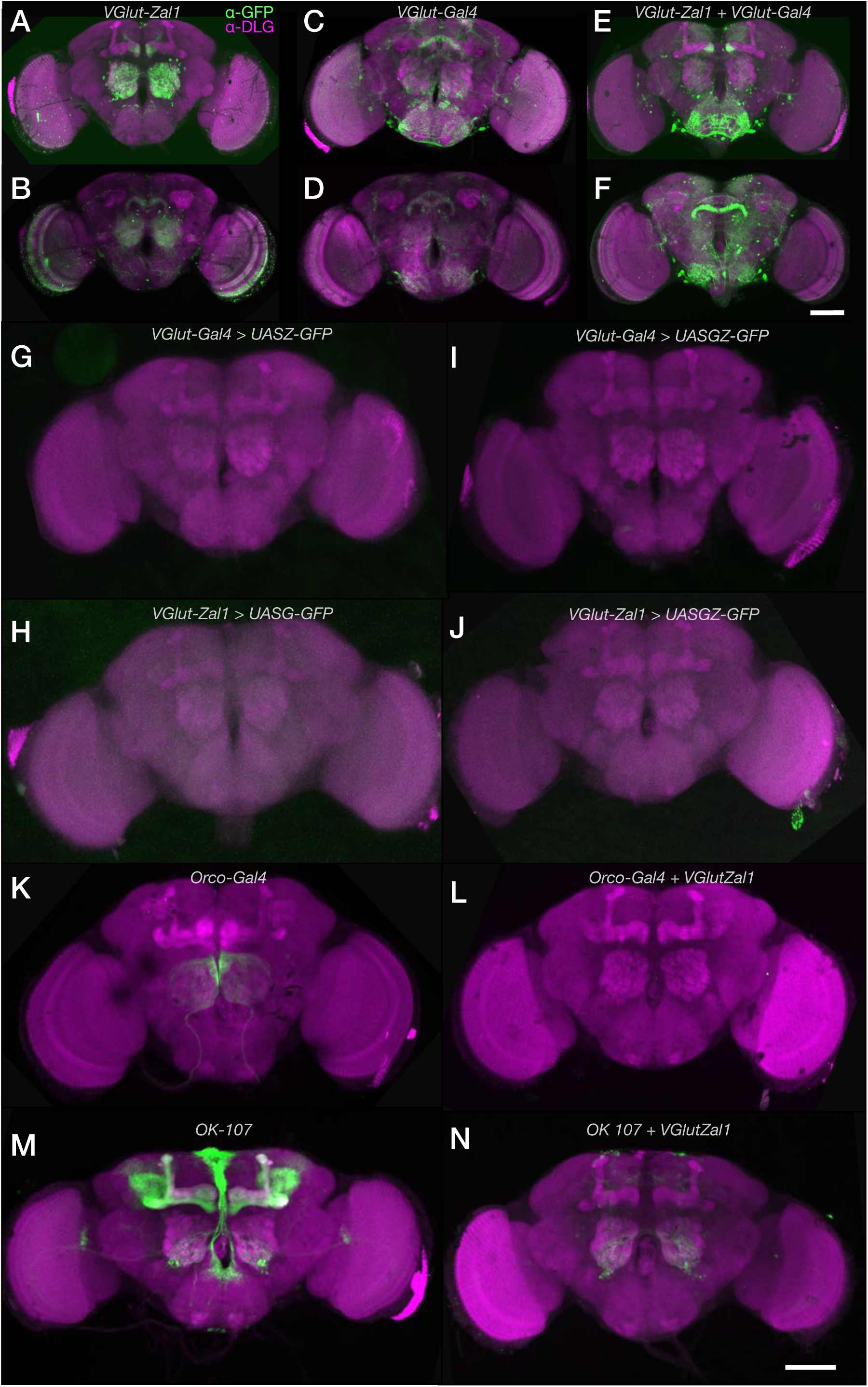
*VGlut-Zal1* drives reporter expression with similar fidelity to *VGlut-Gal4* and generates distinct intersected expression pattern in combination of Gal4 lines. In adult fly brains, widespread GFP expression was observed for both *VGlut* drivers and their combination. **A–B.** Maximum intensity projection images of (A) the brain anterior to the ellipsoid body and (B) the ellipsoid-posterior brain of a *UASZ-mCD8GFP/+; VGlut-Zal1/+* fly stained with α-GFP (green) and α-DLG (magenta) antibodies. **C–D.** Maximum intensity projections of the anterior (C) and posterior (D) expression patterns in a *UASG-mCD8GFP/+; VGlut-Gal4/+* brain stained with α-GFP (green) and α-DLG (magenta) antibodies. **E–F.** Projection images of an anterior (E) and posterior (F) portions of a *UASGZ-mCD8GFP/+; VGlut-Zal1/VGlut-Gal4* brain stained with α-GFP (green) and α-DLG (magenta) antibodies. **G–H.** *VGlut-Gal4* and *VGlut-Zal1* are not individually active at non-cognate UAS sites. *VGlut-Gal4; UASZ-GFP* and *VGlut-Zal1; UASG-GFP* brains stained with α-GFP showed little or no green fluorescence. **I**. *VGlut-Gal4; UASGZ-GFP* and **J.** *VGlut-Zal1; UASGZ-GFP* brains were α-GFP-negative. **K.** Expression pattern of *Orco-Gal4* crossed with *UASG-GFP*. **L.** Intersectional expression pattern of *Orco-Gal4* generated using *VGlut-Zal1;* no GFP expression was observed. **M.** The expression pattern of *OK107-Gal4* crossed with *UASG-GFP*. **N.** The intersectional expression pattern of *OK107-Gal4* with *VGlut-Zal1*. Images show staining with α-GFP (green) and α-DLG (magenta). Scale bars represents 200 μm; dorsal is up.

### Co-expressed Zal1 and Gal4 drive expression from a hybrid UAS

To explore the utility of Zal1 for expression refinement, we made flies carrying *VGlut-Gal4, VGlut-Zal1*, and a responder transgene *UASGZ-GFP*. We hypothesized that a heterodimer of the two transcription factors would drive expression of GFP through the UASGZ hybrid binding sequence. Flies carrying all three transgenes showed GFP expression in many cells, indicating that the Gal4-Zal1 heterodimer did form *in vivo* and was functional at the UASGZ sites (Figure 2E–F). There were qualitative differences between GFP expression in the Zal1-Gal4-UASGZ heterodimer brains and the respective monomer-expressing brains, possibly arising from differences in transgene design. These data verify that Zal1 and Gal4 can activate transcription from a hybrid UASGZ, and thus have the potential to drive expression at an intersection.

### *VGlut* homodimeric lines do not activate non-cognate UAS sites

The specificity of intersectional expression patterns from Zal1-Gal4 combinations is predicated on the specificity of binding to their respective UAS sites: broad cross-reactivity would make an OR operation instead. Possible cross-reactivity was examined in non-cognate UAS/transcription factor controls. A *VGlut-Gal4* line were crossed with a *UASZ-CD8::GFP (‘UASZ-GFP’)* reporter line. Confocal images revealed almost no GFP expression in the brain, indicating that *VGlut-Gal4* by itself does not drive expression from a UASZ responder (Figure 2G). Similarly, a *VGlut-Zal1* line was evaluated by crossing it with *UASG-GFP*; brain expression in the progeny of these crosses was weak (Figure 2H), indicating that cross-reactivity is minimal. As previously reported for their *in vitro* counterparts (Pomerantz et al. 1998), the present results show that *in vivo* Zal1 and Gal4 interact with their cognate UAS sites specifically. To exclude the possibility that homodimeric factors were inappropriately active at the hybrid UASGZ sites, *VGlut-Gal4* flies were crossed with *UASGZ-GFP*. Green fluorescence was low (Figure 2I), indicating that Gal4 activation from tandem UASGZ sites is poor. Similarly, we examined whether *VGlut-Zal1* alone drove robust expression from *UASGZ-GFP* (Figure 2J): it did not.

### VGlut-Zal1 restricts the expression breadth of Gal4 lines

The *VGlut-Gal4*-dependent activity of *VGlut-Zal1* at UASGZ suggested that *VGlut-Zal1* could be useful to restrict the cellular range of existing Gal4 transgenes. To test this idea, we examined enhancer trap lines with and without *VGlut-Zal1*. The *Orco-Gal4* line drives expression in a majority of olfactory receptor neurons (Larsson et al. 2004), sending axonal projections to the antennal lobe (Figure 2K). When *Orco-Gal4* was combined with *VGlut-Zal1* and *UASGZ-GFP*, green fluorescence was absent (Figure 2L). This result likely reflects that *VGlut-Zal1* and the cholinergic olfactory-receptor neurons have no overlap. Another line, *OK107*, drives expression in the mushroom body, the pars intercerebralis and the antennal lobe (Figure 2M). When this line was crossed with glutamatergic Zal1, the mushroom body and pars intercerebralis were absent: only some antennal-lobe cells and a few dorsal cells remained (Figure 2N). The same type of experiment was performed on 16 Gal4 enhancer-trap lines (Hayashi et al. 2002). Compared with these lines’ own generally broad expression ranges, the distributions in combination with *Vglut-Zal1* were sharply more limited (Figure S1A–P). Several of the intersectional brains displayed almost no GFP+ cells *(NP6235, NP2002)*, suggesting that Zal1-Gal4 does not produce broadly mistargeted or ectopic responder expression (Figure S1K’ & Q’).

### Gal4-Zal1 activation is susceptible to Gal80 repression

We aimed to determine whether the Gal4-Zal1 dimer was repressible by Gal80. The *NP4683* enhancer trap line expresses in several areas, including the antennal lobe, mushroom body, the ellipsoid body, subesophageal zone (SEZ) and the ventral nerve cord (VNC) (Figure S2A). As with other lines, *VGlut-ZAL1* intersection produced a reduced expression range; it excluded the mushroom body and antennal lobe expression, but retained GFP in the ellipsoid body, SEZ and VNC (Figure S2B). The *tsh-GAL80* driver represses GAL4 expression in the thoracic and abdominal nervous system (Clyne & Miesenböck 2008). In flies carrying both the ternary system and *tsh-GAL80* (*tsh-Gal80/UASGZ-GFP; VGlut-ZAL1/NP4683*), the ellipsoid body remained brightly GFP+, but the SEZ and VNC expression was diminished (Figure S2C). These data are compatible with the idea that the Gal4-Zal1 dimer is repressible by Gal80.

These qualitative observations show that Zal1-UASGZ is interoperable with both Gal4 and Gal80, and can limit and refine the expression range of existing lines. However, the glutamatergic system is a challenging target for quantitative analyses of expression: the cells are numerous; and the transporter is predominantly present at the nerve terminals—the α-VGLUT antibody labels cell bodies weakly, rendering their identification and quantification inaccessible (data not shown). Therefore, we turned to other neurogenetic systems to quantify Zal1 performance.

**Supplementary Figure S1.**
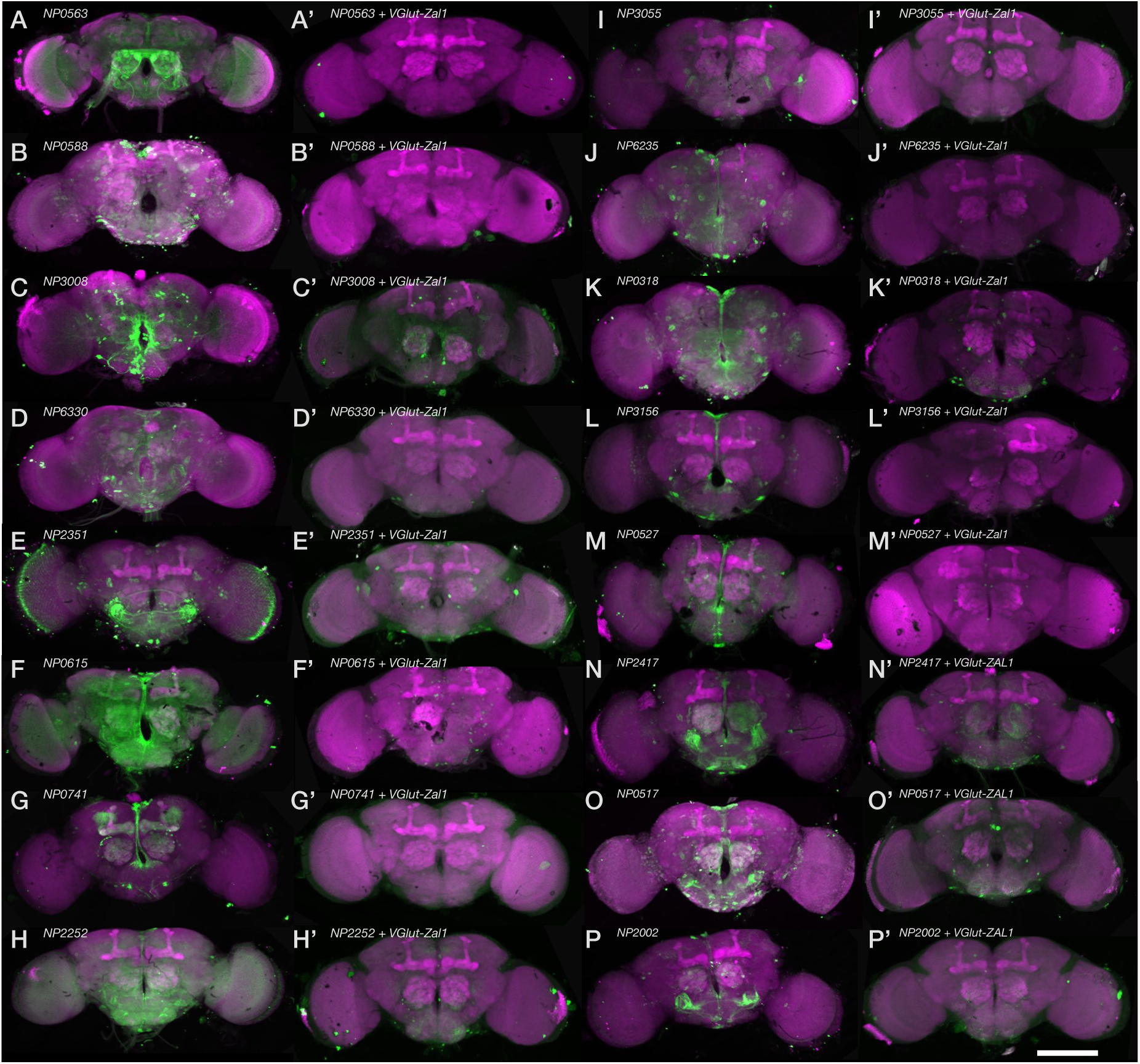
Expression patterns of NP lines with and without *VGlut-Zal1* AND operation. Expression patterns of 16 NP drivers in the adult brain, alone and in combination with *VGlut-Zal1*. All brains are stained with α-GFP (green) and α-DLG (magenta). **A–R.** Expression patterns of NP0563, NP0588, NP3008, NP6330, NP2351, NP0615, NP0741, NP2252, NP3055, NP6235, NP0318, NP3156, NP0527, NP2417, NP0517, and NP2002 Gal4 enhancer trap lines. **A’-R’.** The intersectional expression patterns in combinations with *VGlut-Zal1* are shown in the panels on the right. Several brains lack appreciable α-GFP signal, including NP0563, NP0588, and NP0741. White scale bar represents 200 μm.

**Supplementary Figure S2.**
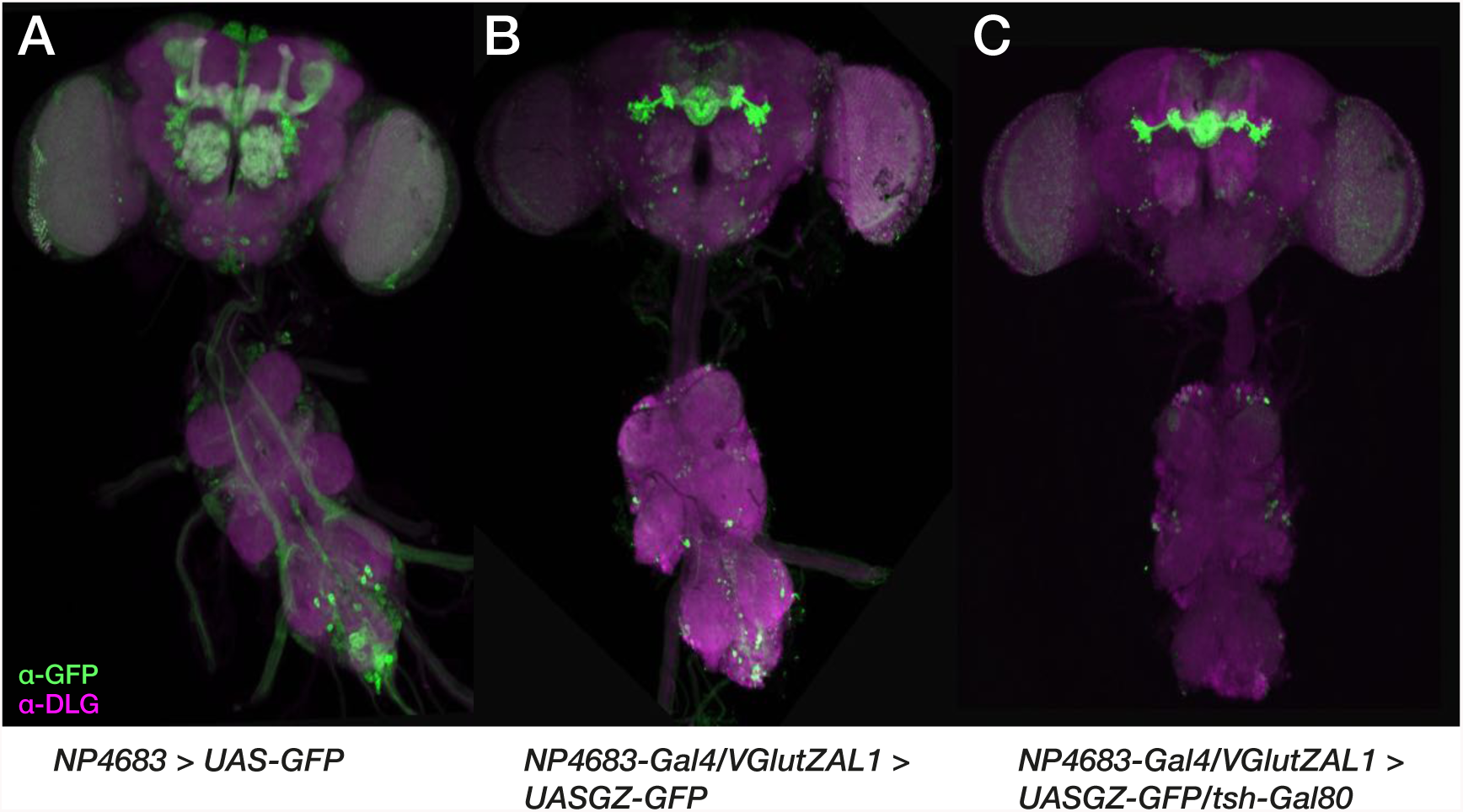
Gal80 represses expression activated by the Gal4-Zal1 dimer. **A.** The enhancer trap line *NP4683* expressed in a wide range of brain cells, as indicated by α-GFP (green) immunostain. The neuropils are stained with α-DLG (magenta). **B.** Intersection with *VGlut-Zal1* reduced the expression range, though left expression in several areas including the ellipsoid body, subesophageal zone, and the ventral nerve cord. **C.** Combination with *tsh-Gal80* left the ellipsoid body brightly stained, while reducing expression in the subesophageal zone, and the ventral nerve cord.

### *Crz-Zal1* drives expression in Corazonergic neurons

To quantify Zal1 performance, we used the Corazonin (Crz) neuropeptide system. The anatomy of these cells is tractable: a *Crz-Gal4* line is available; Crz is expressed in just 6–8 cells per hemisphere; and an α-Crz antibody can be used for Crz^+^ cell identification (Choi et al. 2008). To analyze *Crz-Zal1* brain expression for comparison with *Crz-Gal4*, we fused *Zal1* to the *Crz* enhancer region (Choi et al. 2008). Control brains carrying the non-cognate driver–responder combinations displayed either GFP levels that were undetectable (UASZ, UASG), or weak (UASGZ, Figure 3M–P). This expression in *Crz-Zal1*>*UASGZ-GFP* brains may be due to a mild affinity of Zal1 for the 20 binding half-sites in *UASGZ-GFP*. In cognate, single-driver combinations, both *Crz-Zal1* and *Crz-Gal4* drove strong expression in numerous optic-lobe cells, the ventral nerve cord and in ~7 Corazonergic dorsal protocerebral neurons (Figure 3A–H). This suggested that the two driver types similar patterns. We crossed both drivers with the *UASGZ-GFP* hybrid reporter, and found that the resulting brains had expression patterns nearly identical to the single-driver lines (Figure I–L). Excluding broad ectopic expression in the optic lobes, *Crz-Gal4* has 67% ectopic cells (~15 cells) in the non-optic-lobe brain (Figure 3A–D, see arrow, Figure 4). However, this ectopic expression was excluded when *Crz-Gal4* and *Crz-Zal1* were intersected (Figure 3 J-L, Figure 4), indicating that while both the Zal1 and Gal4 drivers have similar extensiveness within Crz+ cells, *Crz-Zal1* has better fidelity—and establishes that a Zal1 driver can be used to refine a Gal4 driver pattern. These data further verify the hypothesis that Zal1 is useful as an effective intersectional transactivator, and support the idea that Zal1 can be used to improve Gal4 driver fidelity.

**Figure 3.**
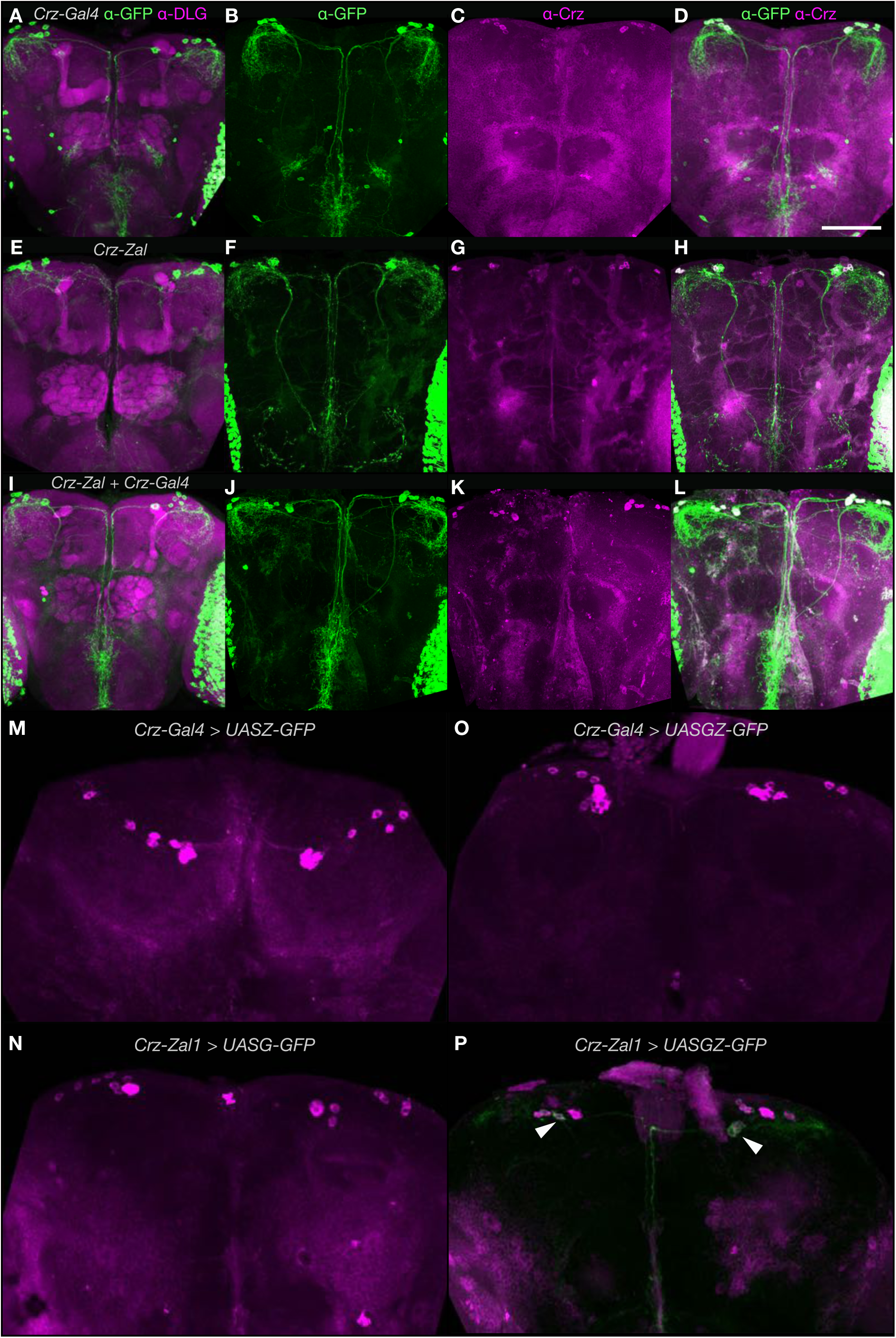
A combination of *Crz-Zal1* and *Crz-Gal4* drives expression in corazonergic cells. Maximum intensity projection (MIP) images of brain immunofluorescence. **A–D.** MIP images of **(A)** of a *UASG-mCD8GFP/+;Crz-Gal4/+* brain stained with α-GFP (green) and α-DLG (magenta) antibodies. **(B)** An image of a *Crz-Gal4/UASZ-mCD8GFP* brain stained with α-GFP (green), **(C)** and α-Crz antibodies (magenta) and **(D)** combined image. **E–H.** *UASZ-mCD8GFP/+; Crz-Zal1/+* brains stained with α-GFP, **(E)** α-DLG (magenta) and **(H)** α-Crz. **I.** *UASGZ-mCD8GFP/+;Crz-Gal4/+* brains stained with α-GFP and α-DLG (magenta) antibodies. **(J)** A *Crz-Gal4/UASGZ-mCD8GFP* brain stained with α-GFP, **(K)** α-Crz, and **(L)** combined image. **M.** Control brains were stained with α-GFP and α-Crz. *Crz-Gal4* is inactive at non-cognate UASZ sites in *Crz-Gal4; UASZ-GFP* brains. **N.** *Crz-Zal1; UASG-GFP* brains stained with α-GFP showed no green fluorescence. **O.** *Crz-Gal4; UASGZ-GFP* brains showed no fluorescence. **P.** *Crz-Zal1; UASGZ-GFP* showed weak expression in a few Crz cells (arrows indicate expression). Scale bar represents 200 μm; dorsal is up.

**Figure 4.**
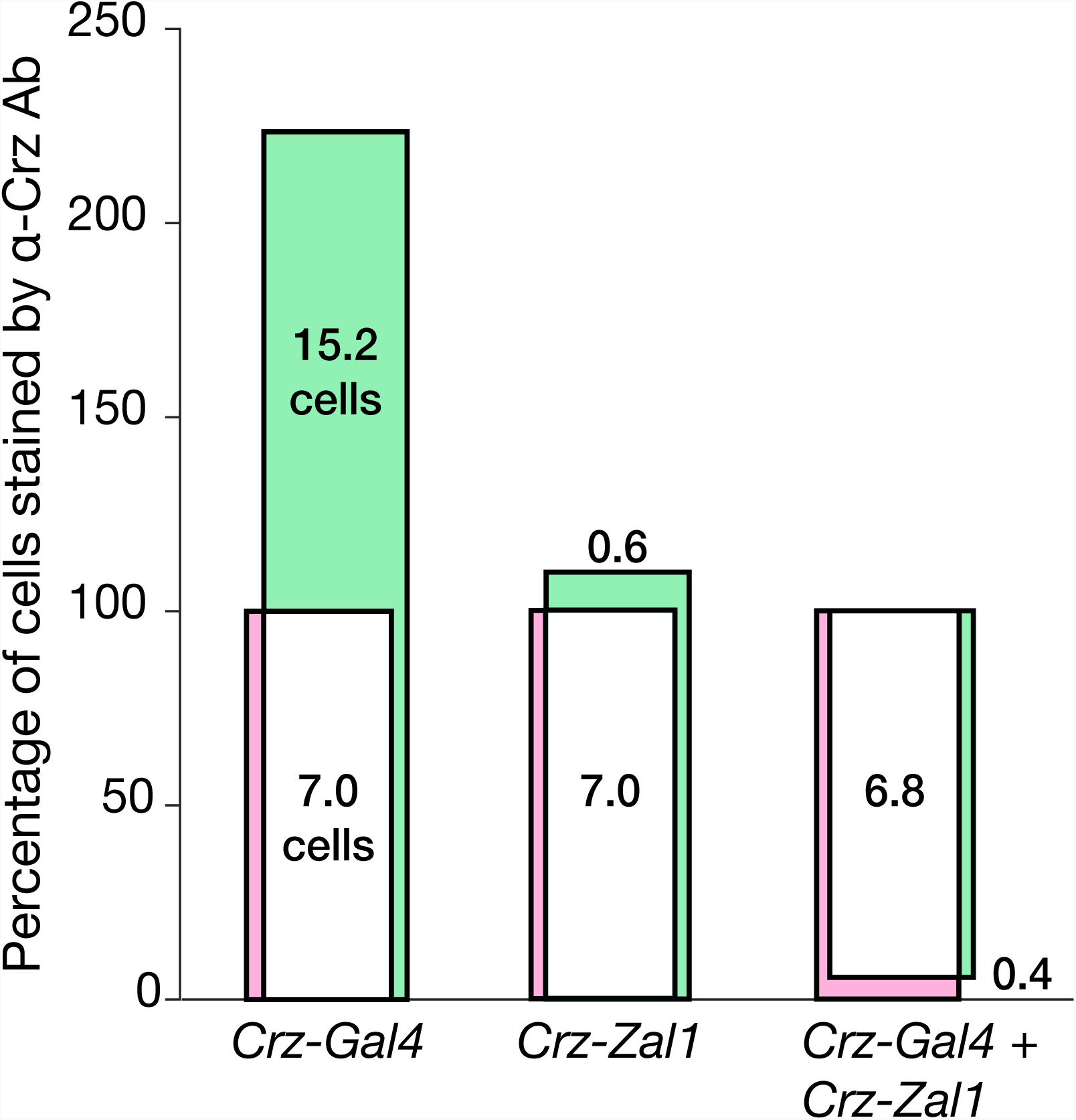
Quantification of Gal4- and Zal1-mediated genetic intersection in Crz cells. A Venn plot shows cell counts of α-GFP and α-Crz antibody staining as percentages. Bar heights are quantitative; bar areas are not. Counts of cells staining positively for Crz were defined as constituting 100% of α-Crz+ cells (magenta bar). Counts of cells staining α-GFP+ were defined as the driver’s expression range (green bar). The overlap between α-Crz+ and α-GFP+ cells is displayed in white. **Left** The *Crz-Gal4* driver expresses GFP in all seven Crz+ neurons, along with expression in 15 ectopic cells. **Center** *Crz-Zal1* expresses in all 7 corazonergic cells along with ectopic expression in less than one cell. **Right** The *Crz-Gal4/Crz-Zal1* double driver expresses in Crz+ cells exclusively.

### *Trh-Zal1* drives expression in serotonergic cells

We tested Zal1 in a third context: the serotonergic system. Serotonin synthesis relies on the *Tryptophan hydroxylase (Trh)* gene; in a Gal4 fusion, the *Trh* enhancer region drives expression in nearly all ~90 serotonergic cells (Alekseyenko et al. 2010). We prepared a *Trh-Zal1* line and assessed expression in controls: *Trh-Gal4* combined with *UASZ-GFP* was inactive; *Trh-Zal1* crossed with *UASG-GFP* had no measurable expression; and green fluorescence in *Trh-Gal4*>*UASGZ-GFP* flies was undetectable (Figure 5M–O). *Trh-Zal1*>*UASGZ-GFP* single-driver brains displayed off-target expression in a few cells, presumably from homodimeric Zal1 activation from the hybrid UASGZ sites (Figure 5P, see arrows). Compared with the controls, the three cognate driver-responder lines revealed expression patterns that were broad and strong. *Trh-Gal4*>*UASG-GFP* expression includes a majority of brain 5-HT^+^ cells (Figure 5A–D): 36 [95CI 32.5, 39.8] cells per hemisphere across nine clusters, with 85.7% fidelity and 90% extensiveness (Figure 6A). Expression in *Trh-Zal1*>*UASZ-GFP* brains was 87.5% extensive, expressing in ~26 serotonergic cells per hemisphere (25.5 [95CI 22, 31]) across five serotonergic clusters (Figure 5E–H), with <2 ectopic cells: 95.7% fidelity (Figure 6A). The double-driver combination *Trh-Gal4*+*Trh-Zal1*>*UASGZ-GFP* expressed in ~24 cells per hemisphere across five cell clusters (Figure 5I–L), representing 82% extensiveness and 100% fidelity (Figure 6A). These results further verify Zal1’s interoperability with Gal4 for intersectional neurogenetics.

**Figure 5.**
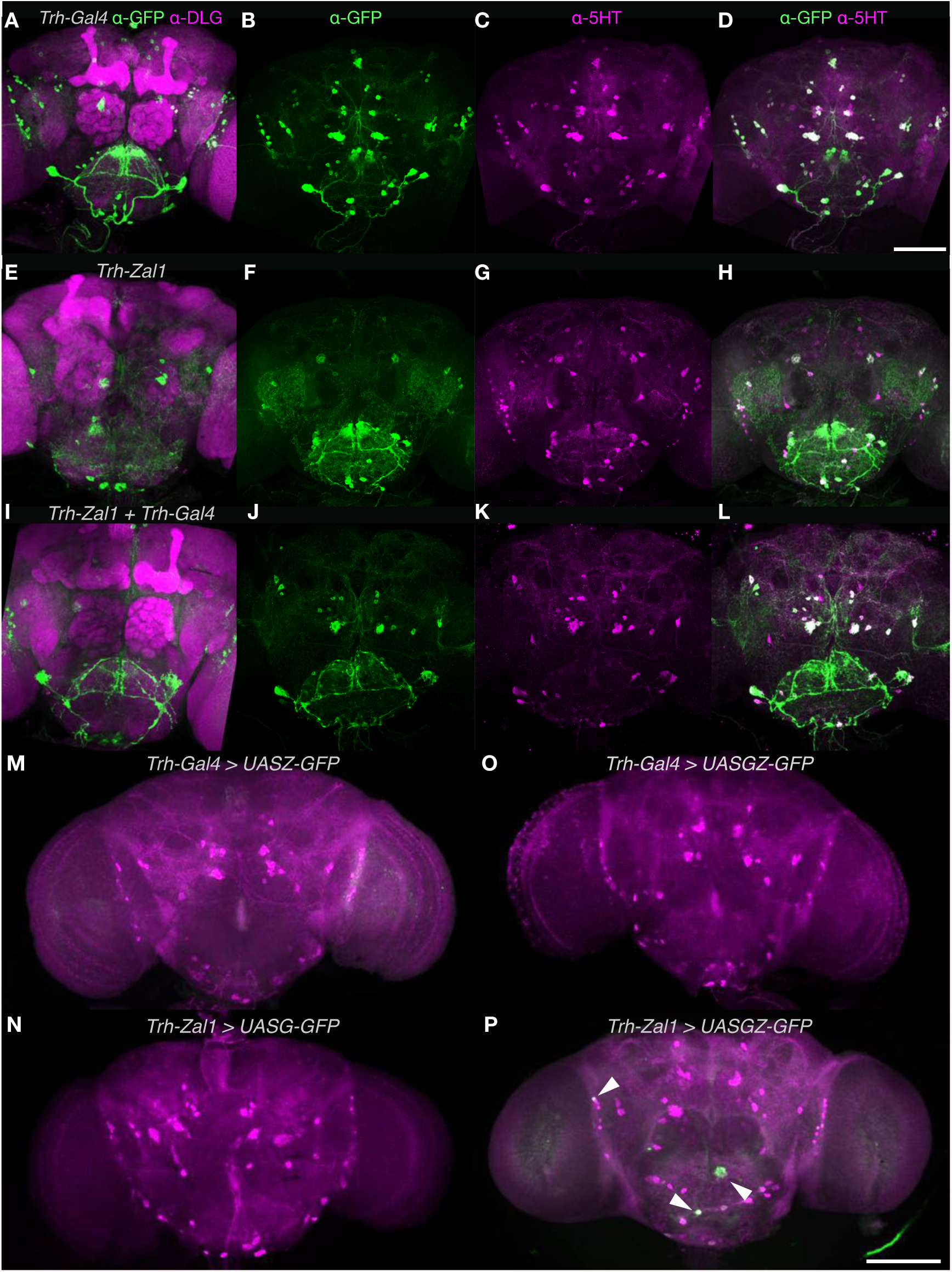
The *Trh-Zal1+Trh-Gal4* combination drives expression in the majority of serotonergic cells. **A–D.** MIPs of **(A)** of a *UASG-mCD8GFP/+; Trh-Gal4/+* brain stained with α-GFP (green) and α-DLG (magenta) antibodies. **(B)** A *Trh-Gal4/UASG-mCD8GFP* brain stained with α-GFP (green), **(C)** with α-5HT antibodies and **(D)** combined image. **E–H.** MIPs of **(E)** a *UASZ-mCD8GFP/+; Trh-Zal1/+* brain stained with α-GFP and α-DLG (magenta); **(F)** a *Trh-Zal1/UASZ-mCD8GFP* brain stained with α-GFP (green), **(G)** with α-5HT, and **(H)** combined image. **I–L.** MIPs of **(I)** a *UASGZ-mCD8GFP/+; Trh-Zal1/Trh-Gal4* brain stained with α-GFP and α-DLG (magenta); **(J)** a *Trh-Zal1; Trh-Gal4/UASGZ-mCD8GFP* brain stained with α-GFP, **(K)** with α-5HT and **(L)** combined image. **M.** A *Trh-Gal4; UASZ-GFP* brain stained with α-GFP showed no green fluorescence **N.** A *Trh-Zal1; UASG-GFP* brain showed no α-GFP fluorescence. **O.** A *Trh-Gal4; UASGZ-GFP* brain showed no α-GFP fluorescence **P.** A *Trh-Zal1; UASGZ-GFP* brain showed weak GFP expression in a few cells; arrows indicate expression. The brains M–P were stained with α-GFP and α-5-HT. Scale bar represents 200 μm; dorsal is up.

**Figure 6.**
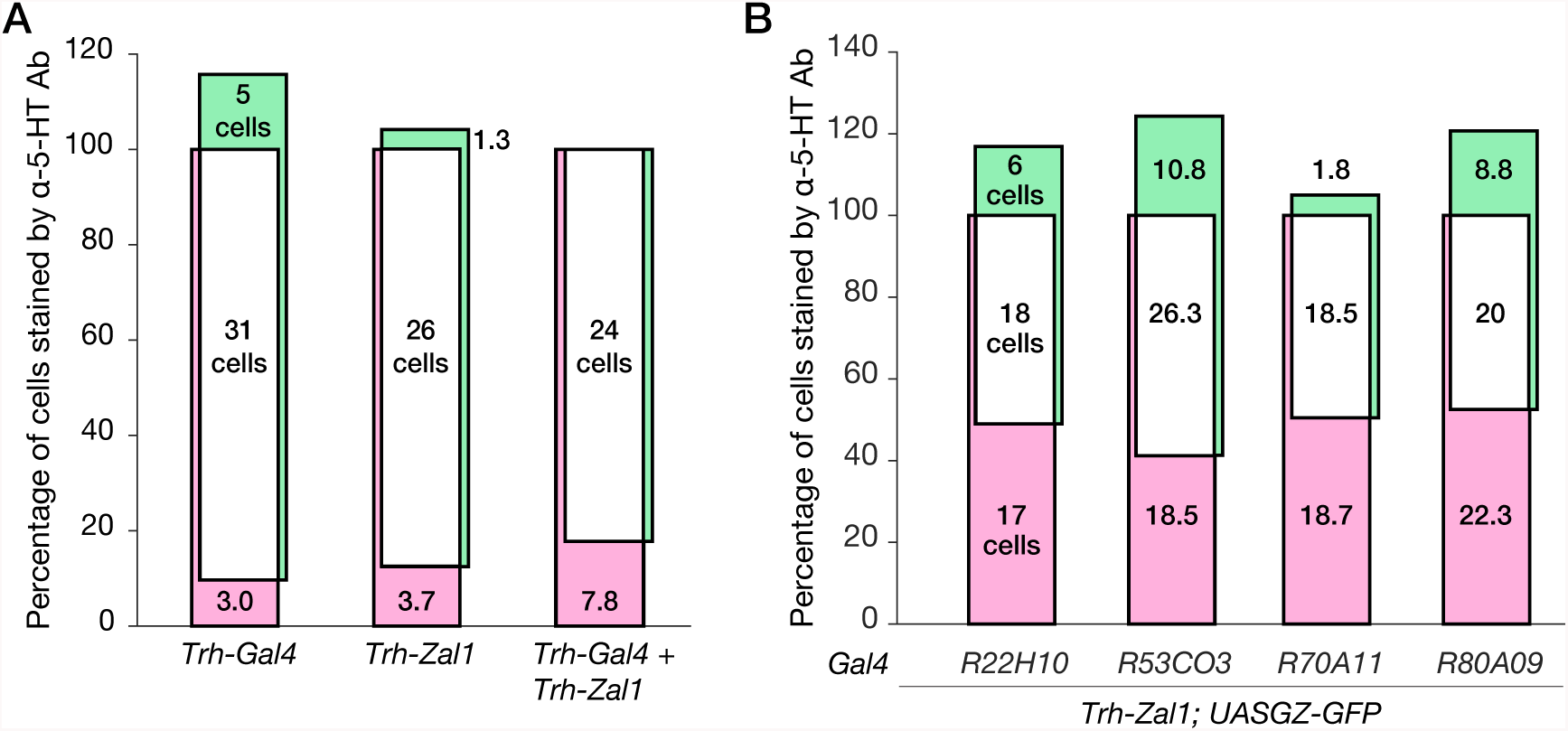
Genetic intersection of Trh-Zal1 with Trh-Gal4 and enhancer trap lines results in high-fidelity expression. **A.** A Venn plot displays α-GFP+ expression as a percentage of α-5-HT+ cells. **Left** *Trh-Gal4* drives expression in 90% of serotonergic neurons, along with 17% of expression in ectopic cells; **Center** similarly, *Trh-Zal1* drives expression in ~88% of serotonergic cells with ectopic expression in 4% of 5-HT+ cells. **Right** The *Trh-Gal4/Trh-Zal1* combination drives expression in ~82% of serotonergic cells with no expression in ectopic cells. The total-count mean of 5-HT^+^ cells ranged from 30 to 34 per brain hemisphere. **B.** The *R22H10-Gal4* + *Trh-Zal1* combination has 51% extensiveness within the antibody stain, with 75% fidelity. The *R53CO3-Gal4* + *Trh-Zal1* combination: 59% extensiveness and 71% fidelity. *R70A11-Gal4* + *Trh-Zal1* combination: 49.5% extensiveness and 91% fidelity. *R89A09-Gal4* + *Trh-Zal1* combination: 47.5% extensiveness and 69.5% fidelity. The total-count mean of 5-HT+ cells ranged from 35 to 42 per brain hemisphere.

### Trh-Zal1–Gal4 combinations improve expression fidelity

While the *Vglut-Zal1* experiments showed that Zal1 can operate with enhancer-trap lines to limit expression, we were not able to quantify the resulting fidelity. Using an α-5-HT antibody that robustly stains fly serotonergic cell bodies, we aimed to test whether *Trh-Zal1* could be used to refine low-fidelity serotonergic Gal4 lines. A visual scan of the FlyLight Gal4 collection (Jenett et al. 2012) found possible serotonergic-driving candidate lines. Subsequent immunostaining of these lines identified four lines expressing in some serotonergic cells (Figure 7A–D). However, these lines included numerous non-5-HT neurons that were densely packed and highly abundant (especially in the optic lobe). Such broad and ectopic expression would confound the interpretation of behavior from such lines; it also prevented quantification of these lines’ serotonin fidelity. Combining these drivers with *Zal1-UASGZ* greatly reduced range while improving fidelity (Figure 7A’–D’). For example, *R22H10-Gal4* drives intense fluorescence in non-serotonergic central-complex cells (Figure 7A); the *Trh-Zal1* AND operation on this driver excluded central-complex expression almost completely. The double-driver combination retained expression in a majority of verified serotonergic neurons: 75% [95CI 71, 80] (Figure 7A’). Overall, counting cells in the four lines found that the mean Gal4-Zal1 intersectional 5-HT+ fidelity was 77% [95CI 65, 92] (Figure 6B). These data verify the hypothesis that Zal1 intersection is useful to refine Gal4 driver specificity.

**Figure 7.**
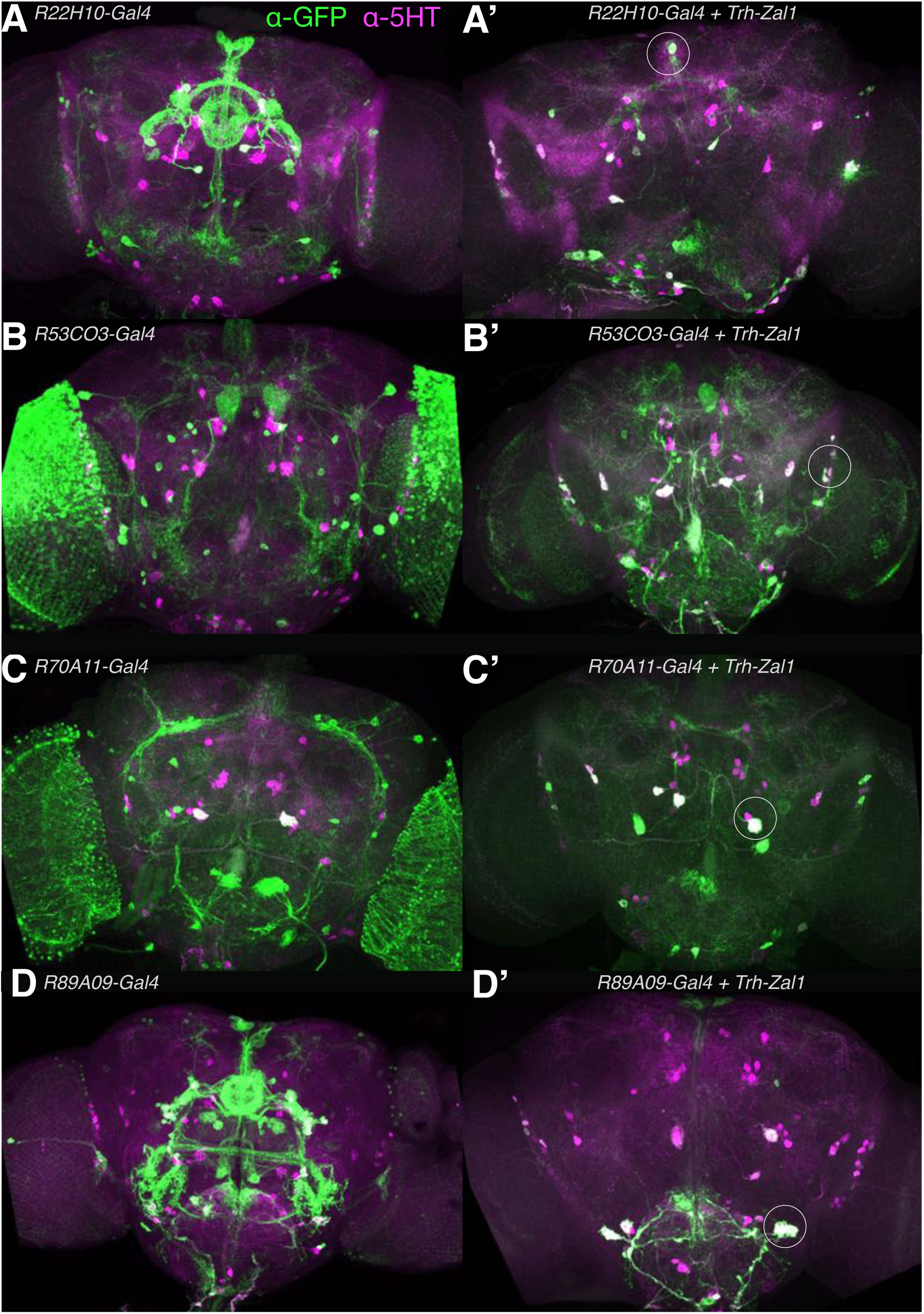
In combination with different Gal4 lines, Trh-Zal1 defines distinct intersectional high-fidelity serotonergic neuronal sets. **A–D.** Expression patterns of *UASG-GFP* signal as driven from Gal4 lines: *R22H10, R53C03, R70A11*, and *R89A09*. Brain images show staining with α-GFP (green) and α-5HT (magenta) antibodies. **A’–D’.** The respective intersectional expression patterns when the drivers are used in combination with *Trh-Zal1*. **A’.** Intersectional expression from *R22H10-Gal4* + *Trh-Zal1* shows highly specific expression in serotonergic LP2 and SE3 cells (arrowhead) **B’.** Intersectional expression of *R53CO3-Gal4* + *Trh-Zal1* shows specific expression in a few LP2, IP and SE1 serotonergic cells. **C’.** Intersectional expression of *R70A11-Gal4* + *Trh-Zal1* shows very specific serotonergic expression: two LP2, two PLP and IP cells. A few ectopic cells can also be seen in the subesophageal zone (SEZ). **D’.** Intersection of *R89A09-Gal4* with *Trh-Zal1* resulted in very specific expression pattern in the serotonergic SE3 cells.

## Discussion

Elucidation of the anatomical and genetic complexity of the brain will require a range of progressively sophisticated tools. Here, we present a method to refine the expression range of existing Gal4 lines. At their non-cognate UAS sites, Gal4 or Zal1 alone have activity that is either weak or absent. Gal4-Zal1 heterodimers are functional at a hybrid *UASGZ* site *in vivo*. Zal1 intersection restricts the number of cells being captured; as judged by antibody counterstaining, Zal1 intersection produces expression with high fidelity.

Other transactivator expression systems have been implemented in *Drosophila*, including LexA-*lexAop* (Lai & Lee 2006; Yagi et al. 2010) and Q (Potter et al. 2010). Such systems can be used with Gal4-UAS in a number of configurations, including driving expression in two distinct cell sets to study their interaction. Our data indicate that *Zal1-UASZ* system can be used this way—in conjunction with Gal4—with a possible benefit of both systems remaining susceptible to Gal80-mediated NOT operations, comparable to intended use of the LexA::GAD fusion protein (Yagi et al. 2010).

The utility of Zal1 can be placed in context with existing Gal4 AND operators. Zal1 is comparable to split-Gal4, though has the added capability of enabling AND operations on the many existing Gal4 lines, a valuable practical benefit. Enhancer trap-driven Flp recombinase can restrict Gal4 expression, though weakly-expressing Flp lines may be affected by stochastic recombination (Sivanantharajah & Zhang 2015). Currently, Gal4 resources include the Kyoto Stock Center’s ~4300 lines, and the Bloomington *Drosophila* Stock Center’s >7000 lines. Many Gal4 drivers capture cellular sets that do not map cleanly to physiological or behavioral function. The ability to restrict expression range with Zal1-UASGZ will further extend the utility of these existing collections.

The Zal1 system has several limitations. First, Zal1 has weak off-target activity at a 20 × UASGZ responder; this means that the combined Gal4-Zal1 expression pattern will include low expression some cells from the Zal1 set. Second, as Gal4-Zal1 is incompatible with existing UASG responder lines, new responder lines for the range of neural inhibitors, activators and other effectors currently will need to be developed. Such efforts, while considerable, will not require the generation of a large collection, will require only a handful of key effectors for most applications, and will augment the utility of the many existing Gal4 lines.

In conclusion, this new expression system provides a versatile tool for the examination of neuronal function, most importantly, for the refinement of Gal4 drivers. Zal1 promises to be a useful method for mapping neural circuits.

## Materials and Methods

### Replacement of the zinc finger in Gal4 with EGR1 domains

A *Gal4* derivative was generated by fusing DNA sequences corresponding to the first two zinc fingers of the mouse transcription activator EGR1 (previously called ZIF268) with DNA coding for residues 41–881 of Gal4, a sequence that includes Gal4’s linker and dimerization domains, as well as the transcriptional-activation regions (Figure 1A–C). Codon-optimized DNA coding for residues 2-59 of EGR1 were synthesized (Genscript Ltd) with an upstream DNA linker that included a KpnI restriction site and 210 base pairs of Gal4 sequence that included an RsrII site. This section was digested and ligated into the pBPGAL4.2Uw-2 vector (Pfeiffer et al. 2010), replacing the first 40 residues of Gal4 while leaving the domains necessary for dimerization and activation intact; this construct was labeled pSVRZal.

### Construction of *VGlut*-, *Trh*-, and *Crz-Zal1* driver lines

To generate drivers that would express Zal1 in glutamatergic cells, serotoninergic, and corazonin (Crz) positive cells, the *VGlut*, *Trh*, and *Crz* enhancer regions were subcloned upstream of *Zal1* to generate *VGlut-Zal1*, Trh-Zal1 and Crz-Zal1 lines respectively. In the case of *VGlut-Zal1*, a 5.5-kb piece of DNA (Daniels et al. 2008) immediately upstream of the *Vglut* translation start site was used. For generating Trh-Zal1 and Crz-Zal1 lines, the same enhancer fragments which have been used to prepare *Trh-Gal4* (Alekseyenko et al. 2010) and *Crz-Gal4* (Choi et al. 2008) were amplified using PCR and subcloned into pSVRZal. For Trh-Zal1, the 1.6 kb promoter region (Alekseyenko et al. 2010) immediately upstream of the *Trh* transcriptional start site was used. For Crz-Zal1, a 434 bp promoter region (Choi et al. 2008) upstream of the putative *Crz* transcription start site was used. All lines were inserted into the *attP2* sites on the 3rd chromosome (BestGene, Inc) of *w^1118^* flies.

### Construction of *UASZ* and *UASGZ* responders

The recognition site of ZFGD1 is a 25-base-pair sequence comprising two inverted six-base-pair EGR1 partial binding sites separated by spacer DNA sequence (Pomerantz et al. 1998). Following the convention set by ‘UASG’, we refer to this palindromic site (AAGCTT-[CGCCCAGAGGACAGTCCTATGGGCGAG × 4]-GACGTC) as ‘UASZ’. Four UASZ sites were introduced using HindIII and AatII sites into the vector pJFRC7-20XUAS-IVS-mCD8::GFP, replacing the original UASG sequences (Pfeiffer et al. 2010) to produce pSVR-4XUASZ-IVS-mCD8::GFP. Four tandem sites were used, as longer repeats of UASZ proved intractable to synthesis subcloning. A non-palindromic, hybrid binding site that combined the recognition half-sites of Gal4 and Zal1 was also synthesized, termed UASGZ (AAGCTT-[CCGGAGTACTGTCCTATGGGCGAG × 20]-GACGTC). To make a GFP responder construct, the UASGZ sites were introduced into the pJFRC7-20XUAS-IVS-mCD8::GFP vector, replacing the original UASG sites using HindIII and AatII sites to generate pSVR-20XUASGZ-IVS-mCD8::GFP. Earlier attempts with 5× UASGZ *Vglut-Zal1* construct produced only weak expression (data not shown). Here synthesis of 20× of tandem sites was successful, an arrangement suitable to maximize expression via the Gal4-Zal1 heterodimer. Both transgenes were targeted to the *attP40* sites on the 2nd chromosome.

### Fly stocks and transgenesis

*Drosophila melanogaster* flies were grown on standard medium at 23°C–25°C. Transgenic animals were generated with the PhiC31-mediated protocol (Bestgene Inc). For brevity, flies transformed with pJFRC7-20XUAS-IVS-mCD8::GFP are referred to as ‘*UASG-GFP*’; flies with pSVR-4XUASZ-IVS-mCD8::GFP are referred to as ‘*UASZ-GFP*’; and flies with pSVR-20XUASGZ-IVS-mCD8::GFP are referred to as ‘*UASGZ-GFP*’. The *VGlut-Gal4* line was a gift from Aaron DiAntonio. *Trh-Gal4* (BL#38389) was procured from the Bloomington stock center. *Crz-Gal4* was a gift from Jae H. Park (The University of Tennessee). The Gal4 lines from the Janelia collection were obtained from Bloomington; NP enhancer trap lines were obtained from the Kyoto Stock Center of the *Drosophila* Genetic Resource Center.

### Immunohistochemistry

Brains were dissected from anesthetized female flies 3–5 days after eclosure and fixed in 4% paraformaldehyde for 30 min at room temperature. Brains were washed for 45–60 min in PBT (phosphate buffered saline with 1% Triton X-100 at pH 7.2). For antibody staining, the samples were further incubated in PBT containing 2% normal goat serum (sc-2043, Santa Cruz Biotechnology) and primary antibodies overnight at 4°C. Primary antibodies were removed by several washing steps (5 × 20 min in PBT) and secondary antibodies were added prior to a second overnight incubation at 4°C). Secondary antibodies were removed with washing in PBT (5 × 20 min) and then finally in PBS (5 × 20 min). Stained brains were mounted in Vectashield (Vector Laboratories, Burlingame, CA, USA) and recorded with confocal microscopy. The following primary and secondary antibodies were used: Alexa Fluor 488 rabbit α-GFP-IgG (A-21311, Molecular Probes, 1:200 dilution), chicken α-GFP (ab13970), rat α-mCD8 (MCD0800, Caltag Laboratories, Chatujak, Bangkok, Thailand), rat α-5-HT (MAB352, Merck), mouse α-DLG1 (4F3 α-DISCS LARGE 1, Developmental Studies Hybridoma Bank, 1:200 dilution), goat α-rat Alexa Fluor 488 (A-11006, Molecular Probes, 1:200 dilution), goat α-mouse Alexa Fluor 568 (A-11004, Molecular Probes, 1:200 dilution), rabbit α-VGLUT (Aaron Diantonio, Washington University, 1:5000 dilution). Rabbit α-Crz (1:500 dilution, Prof. Jan Adrianus Veenstra, University of Bordeaux,France).

### Neuroanatomical comparison of cell sets in NP and GMR lines

With either *UASG-GFP* or in combination with VGlut-Zal1; *UASGZ-GFP*, the following enhancer-trap lines were subjected to α-GFP and α-DLG staining: *Orco-Gal4*, *OK107*, *NP0517*, *NP0588*, *NP3363*, *NP2002*, *NP2417*, *NP3008*, *NP4683*, *NP6235*, *NP6330*, *NP0318*, *NP2351*, *NP3156*, *NP0527*, *NP0615*, *NP0741*, *NP2252*, *NP0563*, *NP3055*, and *NP0564*. With either *UASG-GFP*—or in combination with *Trh-Zal1*; *UASGZ-GFP*—the following GMR module-trap lines were subjected to α-GFP and α-5-HT staining: *R89A09*, *R70A11*, *R53C03*, and *R22H01-Gal4*.

### Microscopy

Serial optical sections were taken in 0.5 μm steps at 1024 × 1024 pixel resolution using a confocal laser scanning microscope (Olympus Fluoview FV1000). Imaging settings for Figure 2 were varied: 28–48% laser power at 488 nm, 39–79% power at 543 nm, 440–744 PMT gain at 488 nm, and 592–794 at 543 nm. Settings for Figure S1 were varied: 20–36% laser power at 488 nm, 29–51% power at 543 nm, 394–614 PMT gain at 488 nm, and 521–708 at 543 nm. Details are provided in a spreadsheet at the Zenodo repository. Settings for Figure 3 were held constant: 5% laser power, 450 PMT gain for the 488 nm laser, and 361 for 555 nm. Settings for Figures 5–7 were held constant: 5% laser power, 537 PMT gain for the 488 nm laser, and 558 for 555 nm. The stacks were visualized and analyzed with the FIJI distribution (www.fiji.sc) of ImageJ (NIH).

### Cell count analysis

For cell quantification, antibody-stained brain samples were scanned in 0.5 μm steps with 1024 × 1024 pixel resolution and 2 μm thickness images were prepared using FIJI to enable counting by eye for all three staining variations - green, magenta, and co-labeled (magenta and green) cell bodies. All cells were counted, including weakly stained cells, so MIP images may appear discrepant from count statistics. Results were reported as the mean count and its confidence intervals; quantification sample sizes were *N_brains_* = 3 samples, *N_hemispheres_* = 6). Venn plots were generated with a custom Matlab script; the vertical axis is meaningful, the bar area is not proportional to the cell count. We used estimation statistics in all cases, reporting means and their confidence intervals (Cumming 2012; Altman et al. 2000; Claridge-Chang & Assam 2016)

### Calculation of driver extensiveness and fidelity

To quantify the quality of different drivers and their combinations, we defined two metrics: *extensiveness* (E) and *fidelity* (F). Extensiveness was measured as how completely a transgenic marker (M+) covers the range of cells identified by an antibody (Ab+) for the cognate protein of the driver’s source gene.

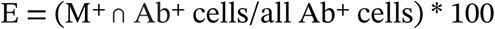

Fidelity was defined as the percentage of marker-positive cells that also immunostained for the cognate protein product.

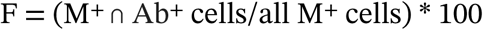

Extensiveness is a desirable property for drivers that aim at capturing a complete set of cells of one neurotransmitter class; however, extensiveness is undesirable for mapping the functions of individual subsets (or individual cells). Fidelity is an unambiguously desirable characteristic.

## Data availability

Microscopy files and other data are available at the Zenodo repository https://doi.org/10.5281/zenodo.1038300.

## Acknowledgements

We thank Aaron DiAntonio for the gift of α-VGlut antibody, Jan Adrianus Veenstra for gifts of α-Crz antibody. We thank Quyen Do for counting cell bodies in image data. We thank Stephen Cohen, Hongyan Wang, members of the Claridge-Chang Lab, and anonymous reviewers for their comments on earlier versions of the manuscript. We thank Yong Zi Tan for the protein model images. We thank Kerry McLaughlin of Insight Editing London for critical review of the manuscript. The α-Discs large 1 4F3 monoclonal antibody was obtained from the Developmental Studies Hybridoma Bank, created by the NICHD of the NIH and maintained at The University of Iowa.

## Author Contributions

*Conceptualization:* ACC; *Methodology:* SVR, CSB, FM, ACC; *Validation:* FM, JYC, SVR; *Investigation:* SVR—*VGlut* experiments, molecular biology, genetic crosses, brain and CNS dissection, immunohistochemistry, confocal laser microscopy, neuroanatomy analysis, FM—*Trh* and *Crz* experiments, molecular biology, genetic crosses, immunohistochemistry, laser microscopy, neuroanatomy analysis, cell counting and analysis, JYC—*Trh* and *Crz* experiments, CNS dissection, immunohistochemistry, laser microscopy, neuroanatomy analysis, CSB—original proof-of-principle experiments, molecular biology, genetic crosses, brain dissection, immunohistochemistry, laser microscopy, JL—NP line screen, immunohistochemistry, laser microscopy, ML—NP line screen, genetic crosses, immunohistochemistry, laser microscopy, SS—molecular biology, genetic crosses; *Writing* – *Original Draft:* SVR; *Writing – Review & Editing:* ACC, FM, SVR; *Visualization:* FM, SVR, JYC, ACC; *Supervision:* ACC—project, SVR & FM—student training and supervision; *Project Administration:* ACC; *Funding acquisition:* ACC.

